# A new sequence logo plot to highlight enrichment and depletion

**DOI:** 10.1101/226597

**Authors:** Kushal K. Dey, Dongyue Xie, Matthew Stephens

## Abstract

**Background:** Sequence logo plots have become a standard graphical tool for visualizing sequence motifs in DNA, RNA or protein sequences. However standard logo plots primarily highlight enrichment of symbols, and may fail to highlight interesting depletions. Current alternatives that try to highlight depletion often produce visually cluttered logos.

**Results:** We introduce a new sequence logo plot, the *EDLogo* plot, that highlights both enrichment and depletion, while minimizing visual clutter. We provide an easy-to-use and highly customizable R package *Logolas* to produce a range of logo plots, including *EDLogo* plots. This software also allows elements in the logo plot to be strings of characters, rather than a single character, extending the range of applications beyond the usual DNA, RNA or protein sequences. We illustrate our methods and software on applications to transcription factor binding site motifs, protein sequence alignments and cancer mutation signature profiles.

**Conclusion:** Our new *EDLogo* plots, and flexible software implementation, can help data analysts visualize both enrichment and depletion of characters (DNA sequence bases, amino acids, etc) across a wide range of applications.

## Introduction

Since their introduction in the early 90’s by Schneider and Stephens [11], sequence logo plots have become widely used for visualizing short conserved patterns known as *sequence motifs*, in multiple alignments of DNA, RNA and protein sequences. At each position in the alignment, the standard logo plot represents the relative frequency of each character (base, amino acid etc) by stacking characters on top of each other, with the height of each character proportional to its relative frequency. The characters are ordered by their relative frequency, and the total height of the stack is determined by the information content of the position. The visualization is so appealing that methods to produce logo plots are now implemented in many software packages (e.g. *seqLogo* [2], *RWebLogo* [17], *ggseqlogo* [18]) and web servers (e.g. *WebLogo* [4], *Seq2Logo* [16], *iceLogo* [3]).

Because the standard logo plot scales the height of each character proportional to its relative frequency, it tends to visually highlight characters that are *enriched*; that is, at higher than expected frequency. In many applications such enrichments may be the main features of interest, and the standard logo plot serves these applications well. However, sometimes it may be equally interesting to identify *depletions*: characters that occur less *often* than expected. The standard logo plot represents strong depletion by the *absence* of a character, which produces less visual emphasis than an enrichment.

To better highlight depletions in amino acid motifs [16] suggest several alternatives to the standard logo plot. The key idea is to explicitly represent depletions using characters that occupy the negative part of the *y* axis. However, we have found that the resulting plots sometimes suffer from visual clutter – too many symbols, which distract from the main patterns of enrichment and depletion.

Here we suggest a simple solution to this problem, producing a new sequence logo plot – the *Enrichment Depletion Logo* or *EDLogo* plot – that highlights both enrichment and depletion, while minimizing visual clutter. In addition, we extend the applicability of logo plots to new settings by i) allowing each “character” in the plot to be an arbitrary alphanumeric string (potentially including user-defined symbols); and ii) allowing a different “alphabet” of permitted strings at each position. All these new features are implemented in our R package, *Logolas*, which can produce generalized string-based logo and *EDLogo* plots. We illustrate the utility of the *EDLogo* plot and the flexibility of the string-based representation through several applications.

## Implementation

### Intuition

In essence, the goal of a logo plot is to represent, at each position along the *x* axis, how a probability vector **p** compares with another probability vector **q**. For example, suppose that at a specific position in a set of aligned DNA sequences, we observe relative frequencies **p** = (*p_A_,p_C_,p_G_,p_T_*) = (0.33,0.33,0.33,0.01) of the four bases {*A, C, G, T*}. The goal of the logo plot might be to represent how **p** compares with the background frequencies of the four bases, which for simplicity we will assume in this example to be equal: **q** = (*q_A_, q_C_, q_G_, q_T_*) = (0.25,0.25, 0.25,0.25). Verbally we could describe the change from **q** to **p** in several ways: we could say “*T* is depleted”, or “*A, C* and *G* are enriched”, or “*T* is depleted, and *A, C* and *G* are enriched”. While all of these are valid statements, the first is the most succinct, and our *EDLogo* plot provides a visual version of that statement. The second statement is more in line with a standard logo representation, and the last is in essence the approach in [16]. See Figure 1.

**Figure 1.**
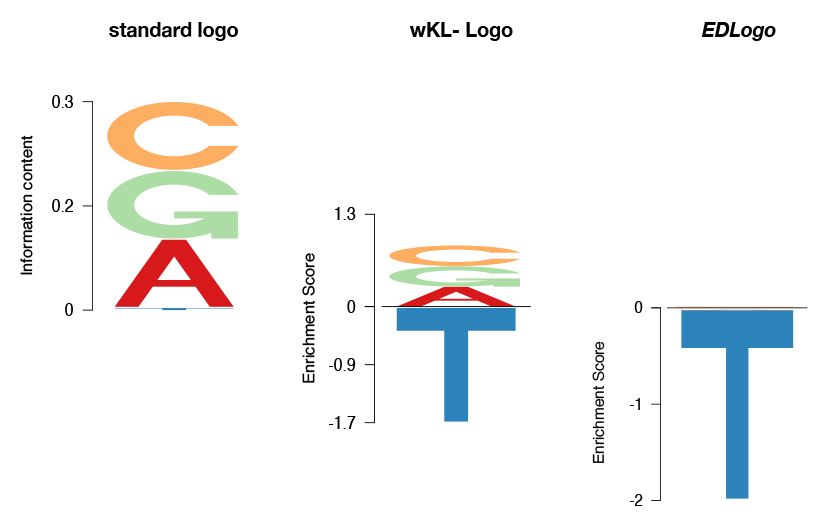
Illustration of the differences between standard logo, *EDLogo* and wKL-Logo representations. The figure shows how the different logos represent observed frequencies **p** = (*p_A_,p_C_,p_G_,p_T_*) = (0.33,0.33,0.33,0.01) (compared with a uniform background, **q** = (0.25,0.25,0.25,0.25)). The standard logo effectively represents **p** by highlighting that “A, C and G are enriched”; *EDLogo* represents it by highlighting “T is depleted”; wKL-Logo represents it as “A, C and G are enriched and T is depleted”. All are correct statements, but the *EDLogo* representation is the most parsimonious.

### The *EDLogo* plot

At a particular position, *j*, of a sequence (or other indexing set), let **p** = (*p*_1_, *p*_2_,…,*p_n_*) denote the probabilities of the *n* elements *C*_1_,…, *C_n_* (which can be characters or strings) permitted at that position, and **q** = (*q*_1_,*q*_2_,…, *q_n_*) denote corresponding background probabilities. Define **r** = (*r*_1_,*r*_2_,…, *r_n_*) by:

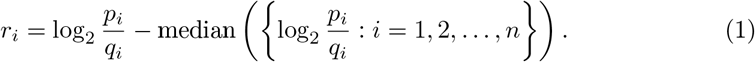

Then at position *j* along the *x* axis, the *EDLogo* plot plots the element *C_i_*, scaled to have height |*r_i_*|, and above the *x* axis if *r_i_* is positive, or below the *x* axis if *r_i_* is negative. Elements are stacked (from bottom to top) in order of increasing *r_i_*, so that the largest characters are furthest from the axis. (In practice, to avoid potential numerical issues if *p_i_* or *q_i_* are very small, we add a small value *ϵ* to each element **p_i_** and *q_i_* before computing *r_i_*; default *ϵ* = 0.01.)

The basic strategy has close connections to ideas in [16], but with the crucial difference that we subtract the median in Equation 1. As our examples will demonstrate, subtracting the median in this way – which can be motivated by a parsimony argument (see below) – can dramatically change the plot, and substantially reduce visual clutter.

Note that the *EDLogo* plot for **p** vs **q** is essentially a mirror (about the *x* axis) of the *EDLogo* plot for **q** vs **p** (e.g. Supplementary Figure S1). We call this the “mirror property”, and it can be interpreted as meaning that the plots treat enrichment and depletion symmetrically. This property is also satisfied by plots in [16], but not by the standard logo plot.

### A model-based view

Suppose we model the relationship of **p** to **q** by

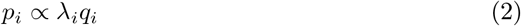

for some unknown (positive) “parameters” λ_*i*_. For example, this model would arise if **q** represents the underlying frequencies of elements in a population, and **p** represents the frequencies of the same elements in a (large) sample from that population, conditional on an event *E* (e.g. a transcription factor binding). Indeed, by Bayes theorem, under this assumption we would have

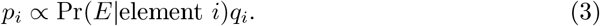

Since the *p_i_* must sum to 1, Σ*_i_pi* = 1, the model (2) implies

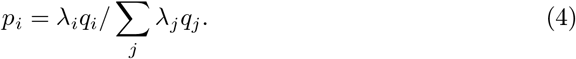

Now consider estimating the parameters λ. Even if **p** and **q** are observed without error, there is a non-identifiability in estimating λ: we can set λ_*i*_ = *cp_i_/q_i_* for any positive *c*. Equivalently, if we consider estimating the logarithms *l_i_*:= log λ_*i*_, we can set

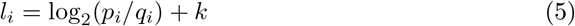

for any constant *k*. Note that *r_i_* in (1) has exactly this form, and so the vector **r** can be interpreted as an estimate of the vector **l**. Furthermore, it is easy to show that, among all estimates of the form (5), **r** has the smallest sum of absolute values. That is, **r** solves the optimization

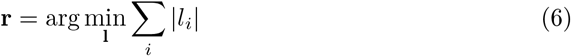

subject to the constraint (5).

Since the sum of absolute values of **r** is the total height of the stacked characters in the *EDLogo* plot, one can think of our choice of **r** as the *estimate of* **l** *that produces the smallest stack of characters* – that is, the most “parsimonious” estimate.

### Interpretation

Roughly speaking, positive values of *r_i_* can be interpreted as indicating characters that are “enriched” and negative values of *r_i_* as indicating characters that are “depleted”. Formally we must add that here enrichment and depletion are to be interpreted as *relative to the median enrichment/depletion across characters*. This relative enrichment does not necessarily imply enrichment or depletion in some “absolute” sense: for example, *r_i_* could be positive even if *p_i_* is smaller than *q_i_*. For compositional data it seems natural that enrichment/depletion be interpreted relative to some “baseline”, and our choice of the median as the baseline is motivated above as providing the most parsimonious plot.

It may also help interpretation to note that for any two characters *i* and *i*′, the difference *r_i_* – *r_i′_* is equal to the log-odds ratio:

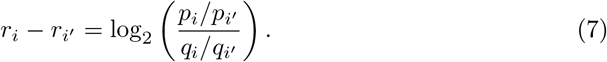

### A variation: the scaled *EDLogo* plot

In the standard logo plot the total height of the stack at each position is scaled to reflect the “information content” at that position, or, more generally, the Kullback-Leibler divergence (KLD) from the background frequencies **q** to the observed frequencies **p** [14]. This scaling highlights locations where **p** differs most strongly from **q**. Similarly, the stack heights in the *EDLogo* plot also reflect the extent to which **p** differs from **q**; for example, if **p** = **q** then the stack height is 0. However, the *EDLogo* stack heights are not equal to the KLD.

Empirically, compared with the standard KLD stack heights, the stack heights in the *EDLogo* plot tend to down-weight locations with a single strongly-enriched element. In settings where this is undesirable, we could avoid it by scaling the *EDLogo* plot to match the standard plot. That is, we could scale the elements at each position by a (position-specific) constant factor so that the stack height is, like the standard plot, equal to the KLD. However, this would lose the mirror property of the *EDLogo* plot because the KLD is not symmetric in **p** and **q**. Thus we instead suggesting scaling by the symmetric KL divergence (symmKLD) between **p** and **q**, which highlights strong single-element enrichments while retaining the mirror property. We call the resulting plot the *scaled EDLogo* plot.

## Results

### Comparison with existing logo plots

Figure 2 illustrates the *EDLogo* plot, and compares it with the standard logo and the weighted Kullback-Leibler logo (wKL-Logo) plot [16], in four diverse applications.

**Figure 2.**
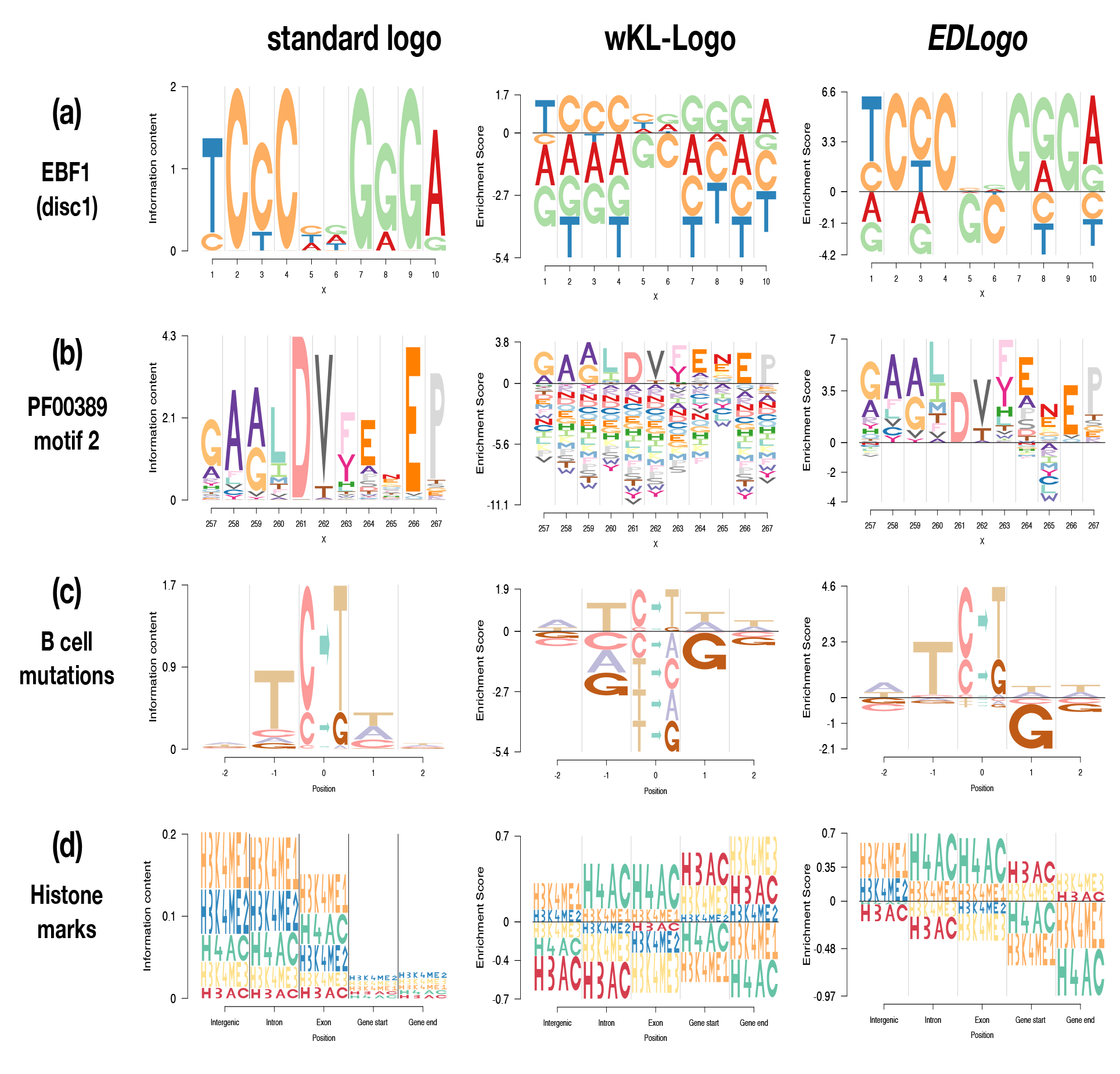
Comparison of standard logo plot, weighted KL (w-KL) logo plot and *EDLogo* plot on four examples. Panel (a): the transcription factor binding site of the EBF1-disc1 transcription factor. Panel (b): the binding motif (Motif2 Start=257 Length=11) of the protein *D-isomer specific 2-hydroxyacid dehydrogenase, catalytic domain (IPR006139)* from [6,12]. Panel (c): mutational signature profile of mutations in lymphoma B cells, with data from [1]. The depletion of G to the right of the mutation – possibly occurring due to the rarity of CpG sites owing to de-amination of methylated cytosines – is clearest in the *EDLogo* representation. Panel (d): relative abundance of histone modification sites across various genomic regions in the lymphoblastoid cell line GM06990 (Table S2 in Koch et al 2007 [8]). These examples illustrate the ability of the *EDLogo* plot to highlight both enrichment and depletion, while avoiding unnecessarily visual clutter. The last two examples also illustrate how our software allows arbitrary strings as elements in a logo plot.

The first two applications (panels (a) and (b)) are settings where the standard logo plot is widely used: visualizing transcription factor binding sites (TFBS) [5,7,9,15,20,21], and protein binding motifs [6,12]. These examples showcase the effectiveness of the standard logo plot in highlighting enrichments: in our opinion it does this better than the other two plots, and in this sense the other plots should be viewed as complementing the standard plot rather than replacing it. These examples also illustrate the differences between the wKL-Logo and *EDLogo* plots, both of which aim to highlight depletion as well as enrichment: the *EDLogo* plot introduces less distracting visual clutter than the wKL-Logo plot, producing a cleaner and more parsimonious visualization that better highlights the primary enrichments and depletions. In particular, for the TFBS example (panel (a), which shows the primary discovered motif *disc1* of Early B cell factor EBF1 from ENCODE [7]), the *EDLogo* plot is most effective at highlighting depletion of bases G and C at the two positions in the middle of the sequence. This depletion is hard to see in the standard logo because of its emphasis on enrichment, and less clear in the wKL-Logo due to visual clutter. This depletion pattern is likely meaningful, rather than a coincidence, since it was also observed in two other previously known motifs of the same transcription factor [5, 9] (see Supplementary Figure S2).

The next two applications (panels (c) and (d) of Figure 2) are non-standard settings that illustrate the use of general strings as “characters” in a logo plot, as well as providing further examples where the *EDLogo* plot is particularly effective at highlighting depletion as well as enrichment.

Panel (c) shows logo plots representing a cancer mutational signature from lymphoma B cell somatic mutations [1]. Here we follow [13] in representing a mutational signature by the frequency of each type of mutation, together with base frequencies at the ±2 flanking bases. We also follow the common convention of orienting the strand so that the mutation is from either a *C* or a *T*, yielding six possible mutation types: *C → T, C → A, C → G, T → A, T → C, T → G*. This Figure panel illustrates two important points. First, it illustrates the flexibility of our software package *Logolas*, which allows arbitrary strings in a logo. For all three logo plots (standard, wKL and ED) we use this to represent the six mutation types by six strings of the form *X → Y*, and we find the resulting plots easier to read than the *pmsignature* plots in [13] (see Supplementary Figure S3 for comparison). Additionally, it also shows that one can use different sets of permitted strings at different positions – strings are only used to represent the mutation in the center, while characters are used to represent the flanking bases. Second, it illustrates a case where, in our opinion, the *EDLogo* plot is a better visual summary than the other plots. Specifically the *EDLogo* plot best highlights the three primary aspects of this signature: enrichment of *C* → *T* and *C* → *G* mutation types; enrichment of *T* at position −1; and depletion of *G* at position +1. Here the depletion of *G* at +1 may be a bi-product of the enrichment of *C* → · mutation types combined with the overall depletion of CpG sites in the genome due to deamination [10]. For readers interested in other cancer mutation signatures, we provide *EDLogo* plots for 24 cancer mutation signatures from [1] in Supplementary Figure S4.

Panel (d) shows logo plots summarizing the *relative* abundance of 5 different histone marks in different genomic contexts (data from lymphoblastoid cell line GM06990, Table S2 (*upper*) of [8]; background probabilities from Table S2 (*lower*) of [8]). Note that relative abundances yield compositional data that can be visualized in a logo plot.

Again this example illustrates the potential to use strings in logo plots. It also represents an example where the *EDLogo* and wKL-Logo plots seem more informative than the standard logo plot. Specifically, the standard logo plot is dominated by the high deviation from background frequencies at the intergenic, exon and intron regions, and the differences in enrichments and depletions among regions are difficult to discern. In comparison, the *EDLogo* and wKL-Logo plots highlight a number of differences among regions (some of which are also noted in [8]). For example, both plots highlight the relative enrichment of H3AC and H3K4me3 near the start and end of genes, and corresponding relative depletion of H4AC and H3K4me1. Both plots also highlight relative enrichment of H3K4me1 compared with other marks in the intergenic, exonic and intronic regions; the relative enrichment of H4AC in intronic and exonic regions, and relative depletion of H3AC in intergenic and intronic regions.

### The *scaled EDLogo* plot

In the first two applications above (panels (a) and (b) of Figure 2) we noted the effectiveness of the standard logo plot in highlighting strong enrichments. This stems from its use of the KLD to scale stack heights at each position. Motivated by this, we implemented a *scaled EDLogo* plot, which combines properties of the *EDLogo* plot (highlighting both enrichments and depletions) and the standard plot (scaling stack heights based on KLD). The *scaled EDLogo* plot for all four of the examples in Figure 2 are shown in Supplementary Figure S5. The results – particularly panels (a) and (b) – illustrate how the *scaled EDLogo* plot tends to emphasize strong enrichments more than the unscaled version, so the scaled version may be preferred in settings where such enrichments are the primary focus.

### Further Variations

Further variations on the *EDLogo* plot can be created by replacing log_2_(*p_i_/q_i_*), in (1) with other functions of (*p_i_, q_i_*), such as the log-odds, log_2_(*p_i_*/(1 – *p_i_*)) – log_2_(*q_i_*/(1 – *q_i_*)). We have not found any particular advantage of such variations over the *EDLogo* plot presented here, but several such variations are implemented in the software and also illustrated in Supplementary Figure S6. In addition, the *EDLogo* strategy of using a median adjustment in (1) to reduce visual clutter can be directly applied to derived quantities such as the position specific scoring matrix (PSSM) [6,12], commonly used to represent protein binding motifs (Supplementary figure S7).

## Discussion

We present a new sequence logo plot, the *EDLogo* plot, designed to highlight both enrichment and depletion of elements at each position in a sequence (or other index set). We have implemented this plot, as well as standard logo plots, in a flexible R package *Logolas*, which offers many other features: the ability to use strings instead of characters; various customizable styles and color palettes; several methods for scaling stack heights; and ease of integrating logo plots with external graphics like ggplot2 [19].

The Logolas R package is currently under active development on Github (https://github.com/kkdey/Logolas). Code for reproducing figures in this paper is available at https://github.com/kkdey/Logolas-paper. Vignettes and a gallery demonstrating features of Logolas are available at (https://github.com/kkdey/Logolas-pages)

## Competing interests

The authors declare that they have no competing interests.

## Author’s contributions

KKD and MS conceived the idea. KKD implemented the package. KKD and DX tested Logolas on the data applications. KKD, DX and MS wrote the manuscript.

## Acknowledgements

We thank Yuichi Shiraishi, John Blischak, Peter Carbonetto, Yang Li and Hussein Al-Asadi for their valuable feedback and helpful discussions. This work was supported in part by NIH BD2K grant CA198933 and NIH grant HG02585.

## Supplementary Methods

Here we detail several alternative options we have implemented for computing the values of *r_i_* when creating an *EDLogo* plot to compare observed relative frequencies **p** with background frequencies **q**:

- *log ratio* approach

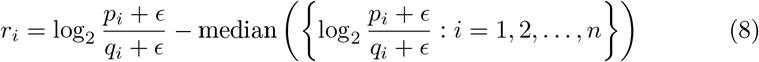
- *log-odds ratio* approach

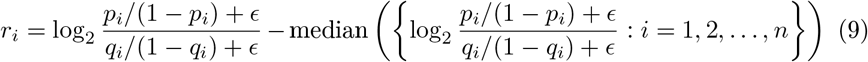
- *ratio* approach

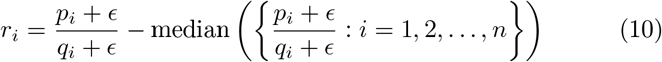
- *probKL* approach [16]

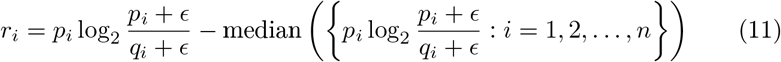

The *log ratio* approach is our default choice and is the one discussed in detail in the main text. Just like the *log ratio* approach, each of the other options also has its corresponding *scaled* version as demonstrated in Supplementary Figure S6.

## Supplementary Figures

**Figure S1.**
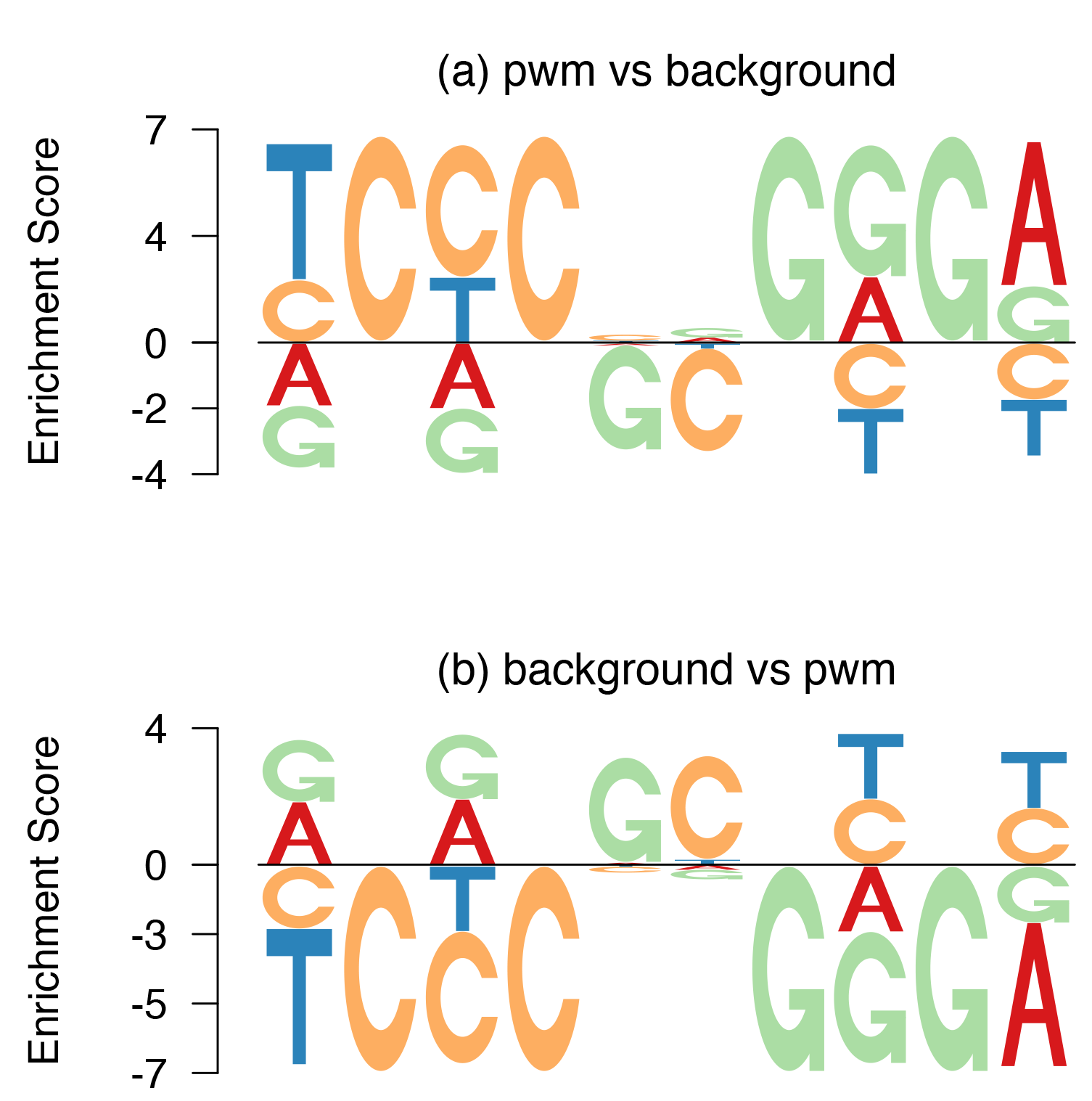
Illustration of “mirror property” of *EDLogo*. *Panel (a): EDLogo* plot of the position weight matrix (PWM) of the primary discovered motif *disc1* from [7] of the EBF1 transcription factor against uniform background. *Panel (b):EDLogo* plot of a uniform PWM against the PWM of EBF1 as background. That is, panels (a) and (b) are comparing the same two PWMs, but differ in which one they treat as the “background”. The *EDLogo* plot obeys the mirror property, in that (b) is a mirror image of (a) (modulo the orientation of the symbols, which are translated and not reflected).

**Figure S2.**
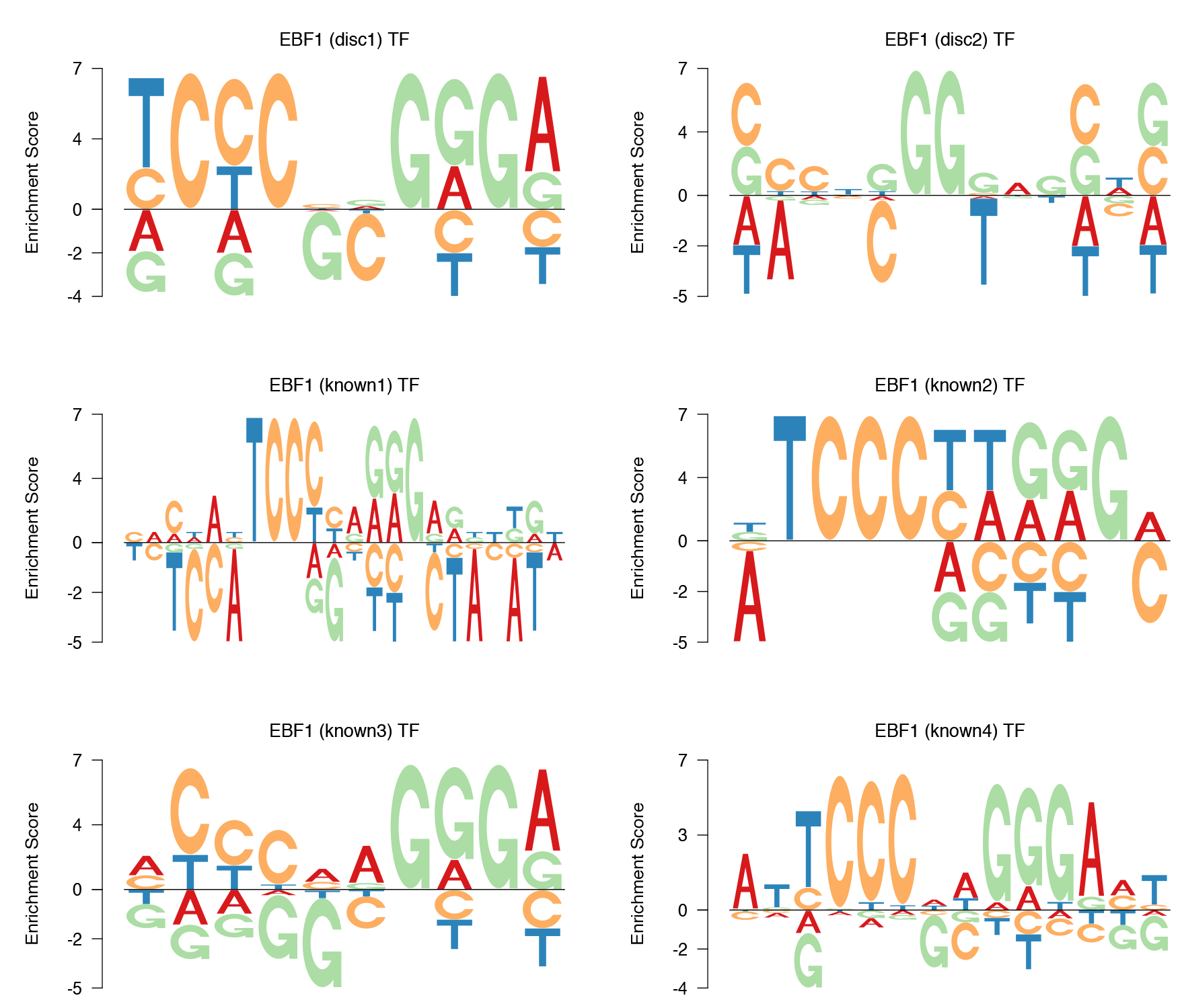
*EDLogo* plots for six different motifs of the EBF1 transcription factor. The PWMS for *known1* and *known2* come from the TRANSFAC database [20]; *known3* from the JASPAR database [9]; *known4* from [5]; *disc1* and *disc2* were discovered by the ENCODE project [7]. Three of the motifs (*known3, known4* and *disc1*) show depletion of G and C in the middle of the binding site.

**Figure S3.**
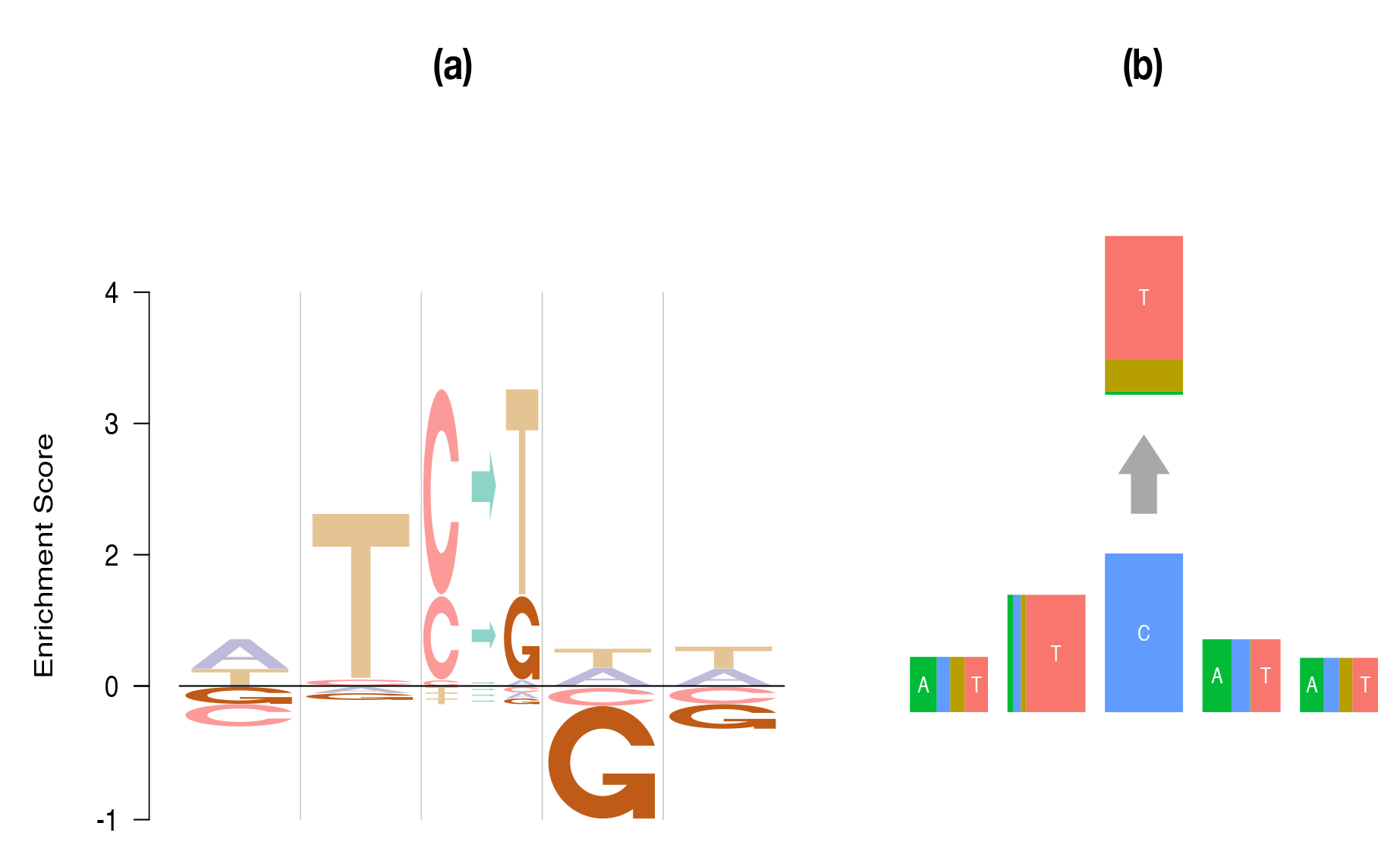
Comparison of the *EDLogo* plot (a) with *pmsignature* [13] plot (b) for visualizing cancer mutational signatures. Both plots show a signature of lymphoma B cell from [1]. The *EDLogo* plot highlights the depletion of G at the right flanking base more clearly than does the *pmsignature* plot. The use of strings to represent mutations in the center is arguably more intuitive than the *pmsignature* representation.

**Figure S4.**
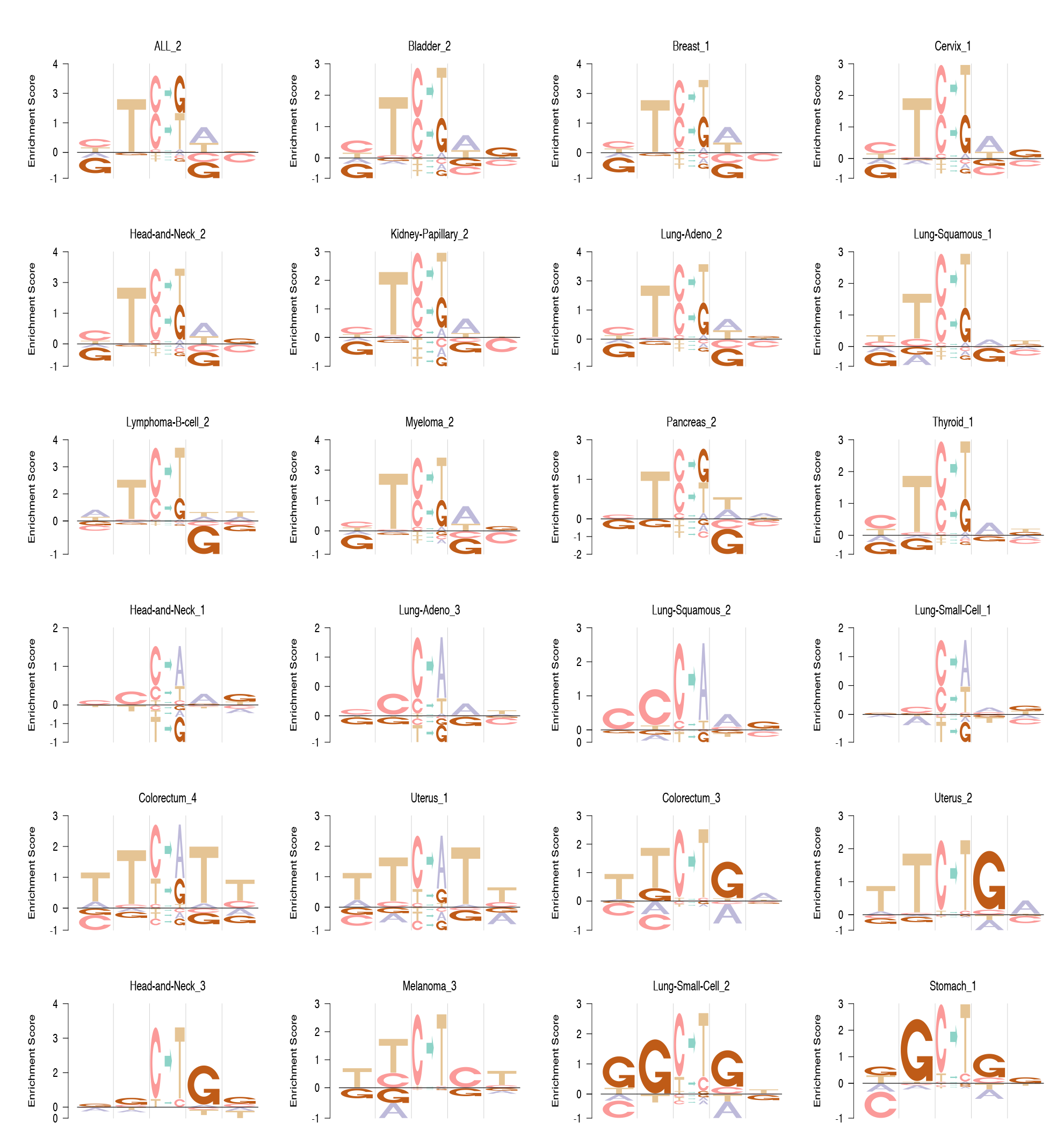
*EDLogo* plots for the mutation signature profiles of 24 different cancer types from [1].

**Figure S5.**
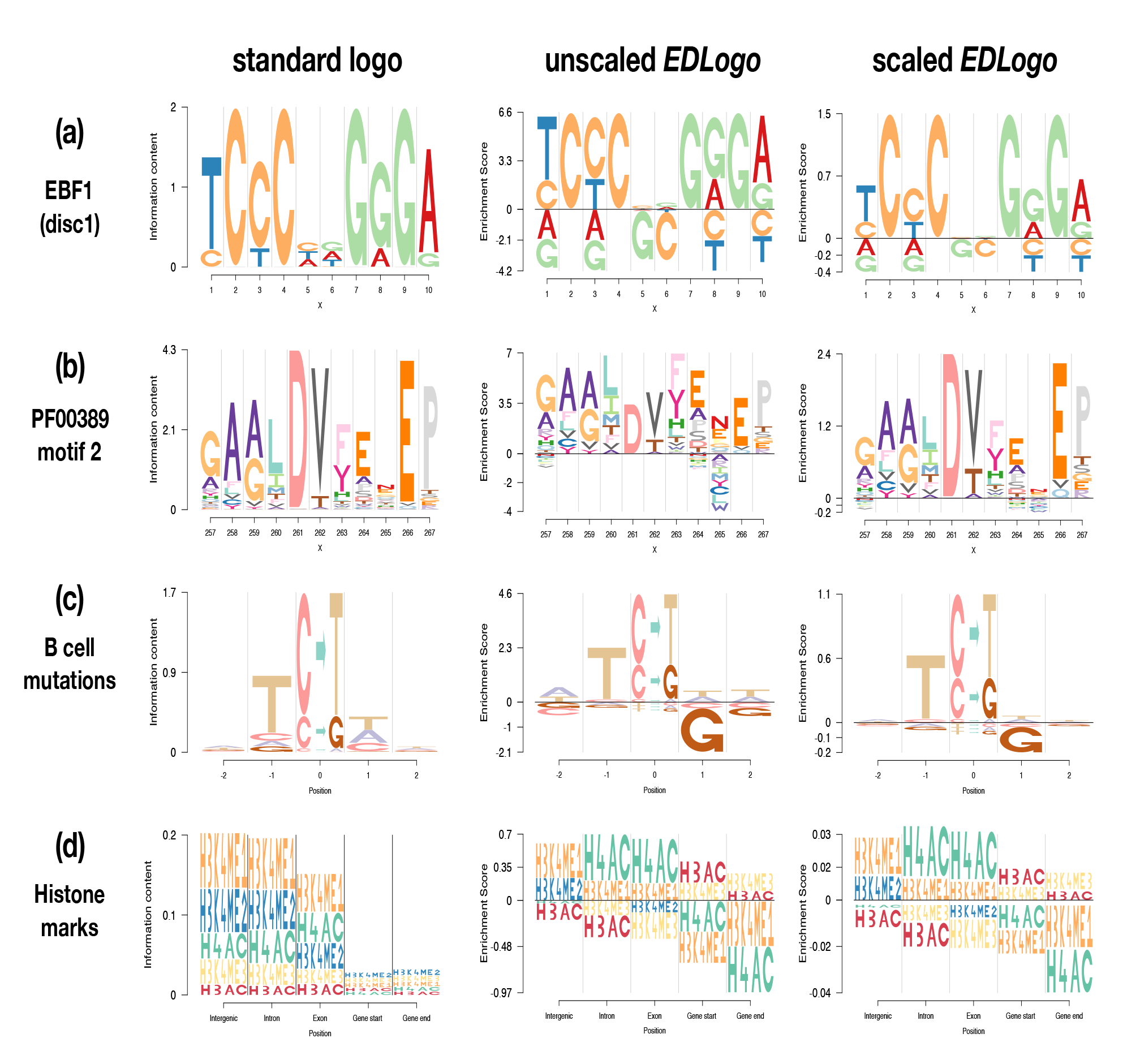
Illustration of *scaled EDLogo* plot on examples from Figure 2. The standard logo and unscaled *EDLogo* plots are repeated here to ease comparisons. The *scaled EDLogo* plot highlights strong enrichments more than the unscaled version and may be preferred in settings when enrichments are the primary focus.

**Figure S6.**
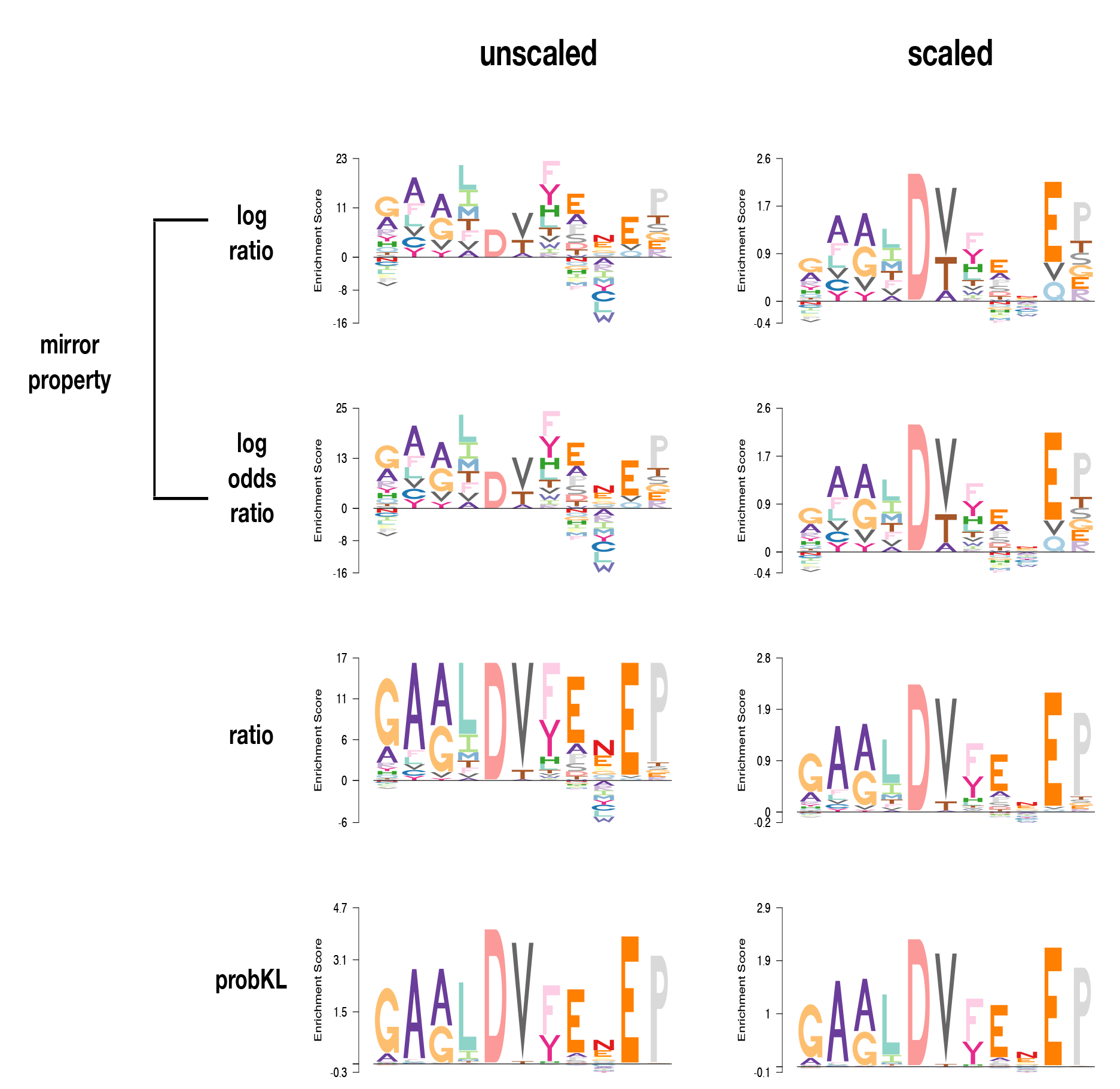
Illustration of various options for *EDLogo* plot. Each plot shows an *EDLogo* plot for a specific binding motif (Motif2 Start=257 Length=11) of the protein *D-isomer specific 2-hydroxyacid dehydrogenase, catalytic domain (IPR006139)* against a uniform background. The plots illustrate the use of several different scoring schemes (*log ratio, log odds ratio, ratio* and *probKL*) with and without scaling by the symmetric Kullback-Leibler divergence. See Supplementary Methods for details on the scoring schemes. (Note that only the *log ratio* and *log odds ratio* scoring schemes satisfy the “mirror property”.)

**Figure S7.**
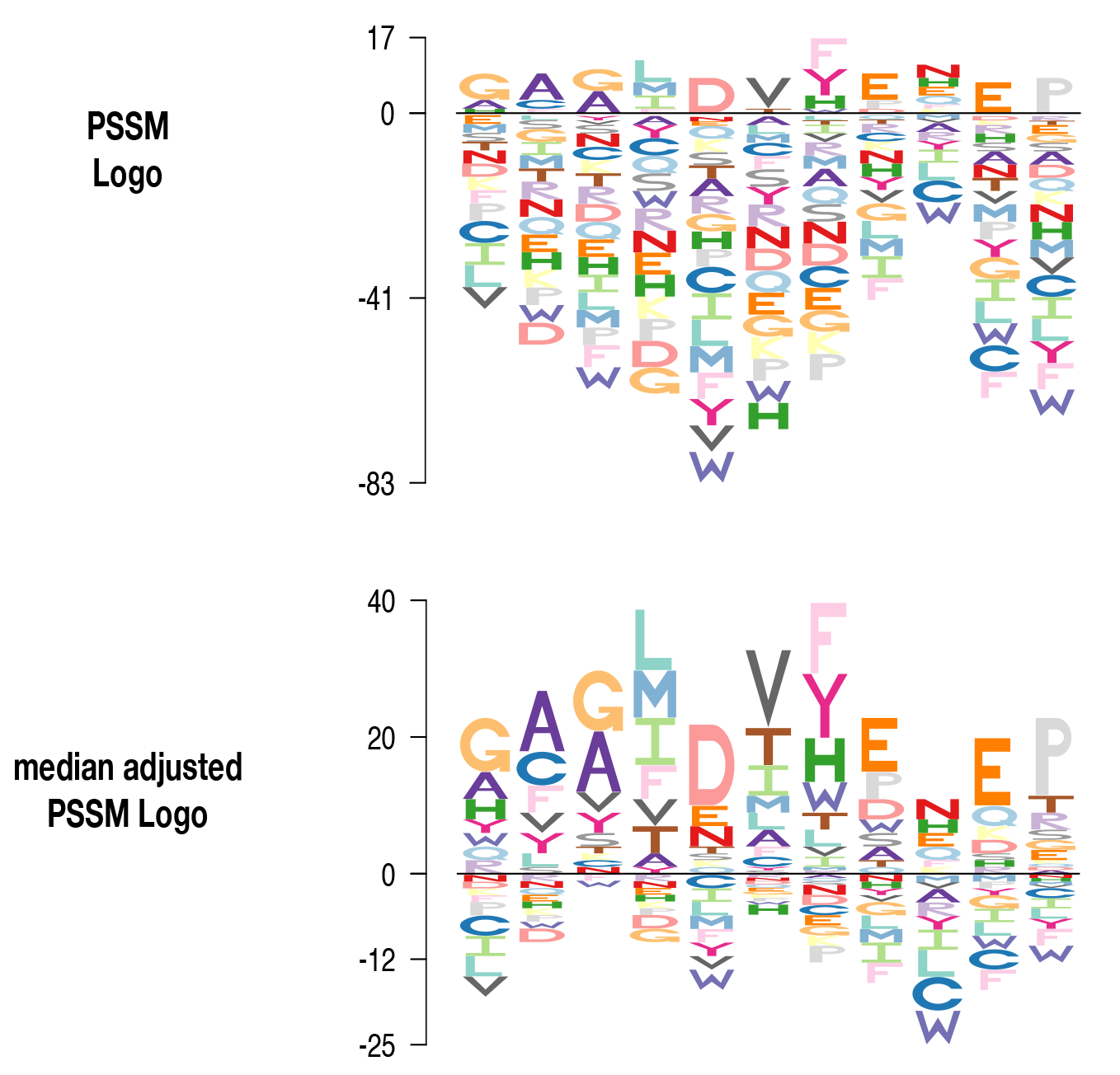
Illustration of median adjustment of a position specific scoring matrix (PSSM). The PSSM shown here is for the binding motif of the protein *D-isomer specific 2-hydroxyacid dehydrogenase, catalytic domain (IPR006139)* (Motif2,Start=257, Length=11). The median adjusted PSSM Logo (*bottom panel*) is arguably less cluttered than the non-adjusted version (*top panel*).

